# Rapid detection of inter-clade recombination in SARS-CoV-2 with Bolotie

**DOI:** 10.1101/2020.09.21.300913

**Authors:** Ales Varabyou, Christopher Pockrandt, Steven L. Salzberg, Mihaela Pertea

**Affiliations:** Center for Computational Biology, Johns Hopkins University, Baltimore, MD; Department of Computer Science, Johns Hopkins University, Baltimore, MD; Department of Biomedical Engineering, Johns Hopkins University, Baltimore, MD; Department of Biostatistics, Johns Hopkins University, Baltimore, MD

**Keywords:** SARS-CoV-2, COVID-19, recombination, coronavirus

## Abstract

The ability to detect recombination in pathogen genomes is crucial to the accuracy of phylogenetic analysis and consequently to forecasting the spread of infectious diseases and to developing therapeutics and public health policies. However, previous methods for detecting recombination and reassortment events cannot handle the computational requirements of analyzing tens of thousands of genomes, a scenario that has now emerged in the effort to track the spread of the SARS-CoV-2 virus. Furthermore, the low divergence of near-identical genomes sequenced in short periods of time presents a statistical challenge not addressed by available methods. In this work we present Bolotie, an efficient method designed to detect recombination and reassortment events between clades of viral genomes. We applied our method to a large collection of SARS-CoV-2 genomes and discovered hundreds of isolates that are likely of a recombinant origin. In cases where raw sequencing data was available, we were able to rule out the possibility that these samples represented co-infections by analyzing the underlying sequence reads. Our findings further show that several recombinants appear to have persisted in the population.

## Introduction

Since the beginning of 2020, the COVID-19 pandemic caused by a newly emerged strain of a betacoronavirus, SARS-CoV-2, has been responsible for over 950,000 deaths and over 30 million infections to date (Dong et al., 2020). The strain has been hypothesized to have emerged as a result of a recombination event between a bat and a pangolin (Zhang et al., 2020), although its precise origin is not yet known. To date, the genetic diversity of the SARS-CoV-2 has been increasing slowly compared to other RNA viruses, with 5 to 7 major circulating clades being identified based on multiple variants common to large numbers of isolates in the GISAID database (Hadfield et al., 2018; Shu & McCauley, 2017). This relative stability of the genetic content of the circulating forms of the virus is promising for the development of vaccines and therapeutics, as well as general understanding of the biology and pathology of SARS-CoV-2.

However, as with other RNA viruses, coronaviruses are known to undergo mutations at high rates (Drake & Holland, 1999). Inter- and intra-host recombinations are also well-studied and occur frequently (Su et al., 2016). As more mutations and lineages of SARS-CoV-2 get fixed in the population, a recombination event caused by a co-infection of a single patient with particles of distinct clades may lead to emergence of novel lineages, posing risks to the efficacy of future vaccines. In fact, several accounts of recombination events in SARS-CoV-2 have been reported in recent months (VanInsberghe et al., 2020; Yi, 2020). As such, rapid and consistent surveillance of the sequenced genomes of SARS-CoV-2 for both novel mutations and recombinations is critical to the development of effective treatments and vaccines (Demir et al., 2020).

Multiple computational methods have been developed to detect recombination in microbial genomes and have been used in studies of HIV-1 mutagenesis, bacterial evolution, and other applications (Posada, 2002). Some popular methods, such as 3seq, analyze every possible triplet of a set of genomes and statistically evaluate the similarity of a sliding window across the query genome to the other sequences of the triplet (Lam et al., 2018). Other methods, like PhiPack, are designed to work for low-divergence genomes but still require a significant number of variants (1-5%) to perform statistical analysis (Bruen et al., 2006). A limitation of these methods for large-scale surveillance is that the algorithms are computationally intensive and require significant resources and time to perform analysis, particularly for the amount of data (nearly 100,000 genomes to date) that has been generated for SARS-CoV-2. There exist 512 trillion unique triplets of sequences for the currently available SARS-CoV-2 genomes, and a similarity analysis for each triplet is computationally infeasible. A more efficient approach is necessary if we want to be able to detect recombinants in a realistic amount of time.

In this work we present Bolotie, a new algorithm designed to conduct mutational analysis and to detect recombinant forms and other anomalies among a very large set of viral sequences. The methods presented are also designed such that novel sequences can be analyzed efficiently without the need to rerun the entire protocol.

We applied Bolotie to search for recombination events in 87,695 complete genomes of SARS-CoV-2 currently available in the GISAID database. In our analysis we identified multiple unique cases of recombination between 4 prominent clades of the virus. Several of the identified recombination events, including some previously reported (VanInsberghe et al., 2020), appear in multiple isolates, suggesting transmission in the population.

Lastly, we propose a methodology for distinguishing true recombination events from cases of mis-assembly of isolates from a host co-infected with several distinct lineages of a pathogen. The proposed method can be applied by to verify future SARS-CoV-2 genomes prior to database submission.

## Results

We first aligned all genomes to the Wuhan-Hu-1 reference genome (GenBank accession <u>MN908947</u>), from which we detected 84,322 single-nucleotide variants (SNVs) at 29,503 sites. After removing all variants that appear in fewer than 100 sequences, we retained a set of 659 SNVs at 411 unique sites. Alignments also revealed 1,349 unique structural variants (934 deletions and 415 insertions). While 2 deletions and 1 insertion were present in over 100 genomes, for the purpose of computational efficiency we did not consider them further.

Using Bolotie on the set of well-supported variants to search for recombination events between sequences in the 4 major clades of SARS-CoV-2, we identified 225 possibly recombinant genomes. Figure 1 illustrates that many of the identified recombination events were represented by a single genome. However, several lineages with near-identical sequences and the same breakpoint sites were observed. In Figure 1 these lineages appear as broad red bands with a high density of outgoing arcs. Additionally, several smaller groups of near-identical recombinant signatures have been observed in which genomes differed by one or two variants. Those genomes were often found to be neighbors in the computed maximum likelihood (ML) trees and often had the same or neighboring inferred parental sequences.

**Figure 1.**
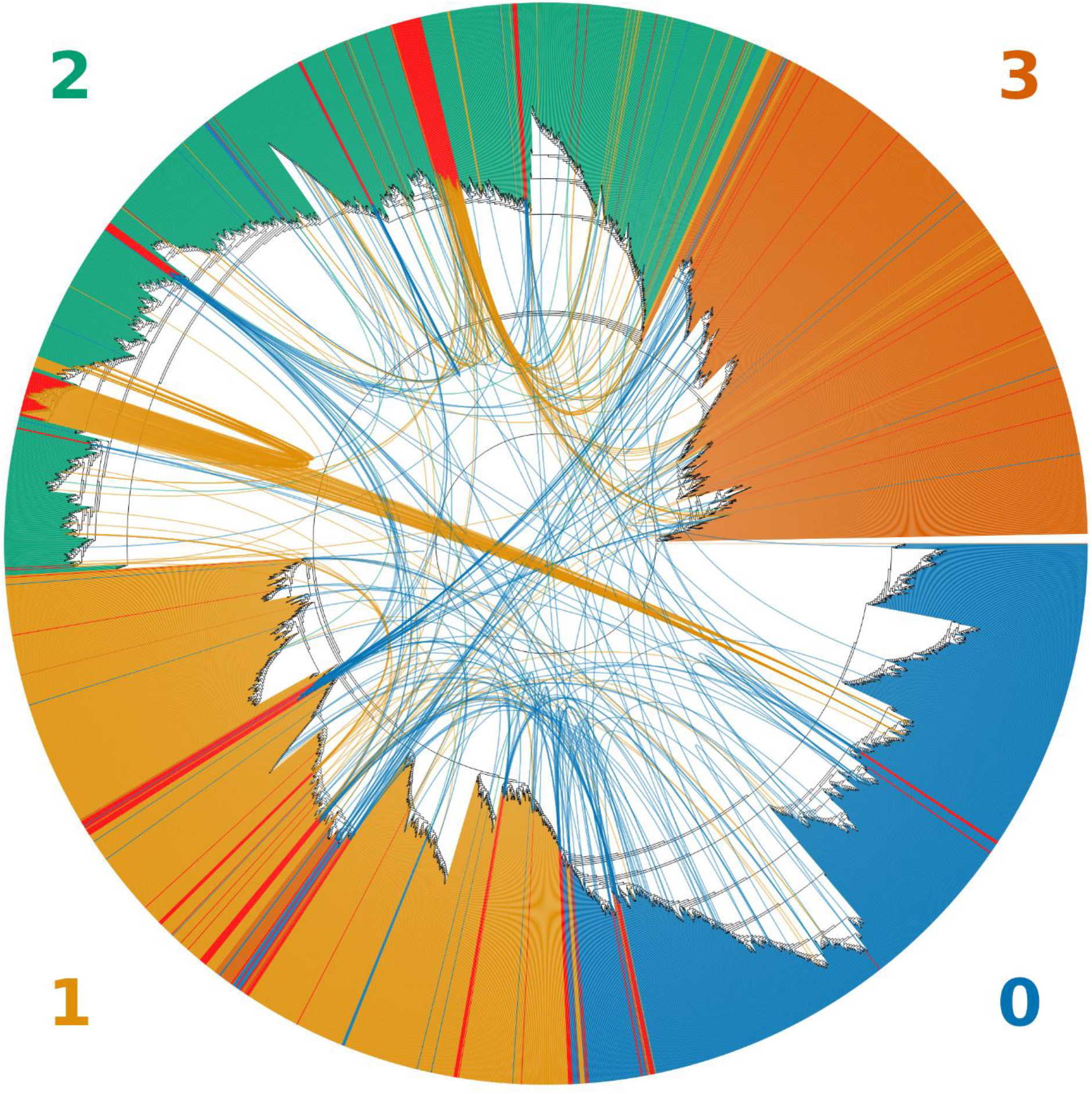
An unrooted topological cladogram of 4,249 SARS-CoV-2 genomes including 225 recombinants labeled as red bars. Arcs link each recombinant to both inferred parental genomes. The color of the arc corresponds to the color of the clade to which a recombinant was clustered within the tree. Clades correspond to the GISAID clades GR (**0**), GH (**1**), G (**2**) and all minor lineages combined (**4**).

Of the 225 recombinant genomes, 109 were labeled in clade #0, 111 in clade #1 and 5 in clade #2 (**Figure 1**). Recombination events happened between members of all 4 clades, with 171 parental genomes identified in clade #0, 41 parental genomes in clade #1, 148 in clade #2 and 90 in clade #3. Additionally, 15 out of 225 potential recombinants were found present in the set of 4,039 representative genomes used by NextStrain (Supplementary Table 2).

Of the 225 identified recombinants, a majority of the recombinant signatures had 1 or 2 breakpoints like the ones shown in Figure 2A, 2B and 2C. However, at least 6 genomes including the one depicted in Figure 2D exhibited more complex patterns of mosaicism with 3 breakpoints.

**Figure 2.**
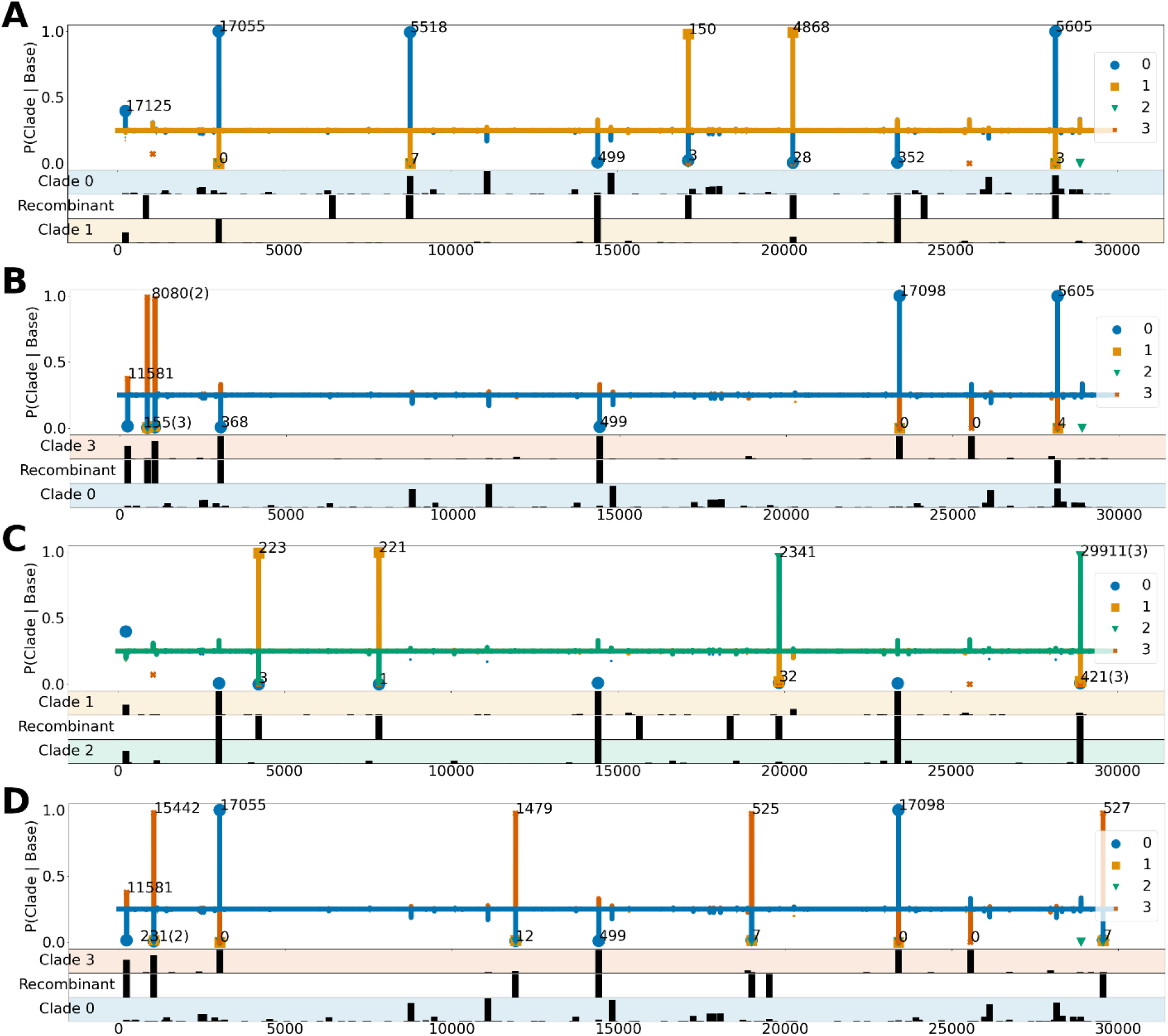
Four examples of inferred recombinant sequences: A. EPI_ISL_439137; B. EPI_ISL_468407; C. EPI_ISL_509874; D. EPI_ISL_417420. The top section of each plot shows conditional probabilities of a clade given a nucleotide at each position. Bars are plotted for the two parent clades and the other clades are shown as dots of the corresponding color. Each peak >0.1 above the baseline (0.25) is labeled with the number of genomes it appears in. An average is reported whenever there are multiple variants in close proximity on the plot, listing the number of averaged variants in parentheses. The three lower panels of each plot show the frequency of variants at each position for parental clades (top and bottom rows) and variants observed on the recombinant genome (middle row).

While in majority of cases the path-finding algorithm of Bolotie relied on clade-defining variants with high conditional probabilities > 0.9, several positions exhibited an inverse pattern and were also useful in the analysis. For example, in Figure 2B (further detailed in Table 1), mutations of cytosine (C) to thymine (T) at position 14,407 and adenine (A) to guanine (G) at position 23,402 are not characteristic of clade 1 (yellow) since they have the same conditional probability of ~0.33 of defining clades 2 and 3. However, these positions are informative in an inverse way, namely that observing a C and an A at those positions is very unlikely if the sequence originated in clade 0. This adds additional evidence to the recombinant origins of the genomes. Similar instances in other genomes aided Bolotie in the search. In fact, since all 4 of the plots in Figure 2 describe recombinants which involve clade 0, the same conditional probabilities at position 14,407 can be observed in each subfigure.

**Table 1.** Mutational signature of the EPI_ISL_439137 recombinant isolate (Figure 2B). The table shows all positions with defining conditional probabilities for each of the parental clades. Read counts extracted from the data deposited in EBI are provided to illustrate the likely single-isolate origin of the genome. Positions are colored according to the respective clade colors used in the manuscript.

| Position | Reference | Observed | P(0\|Base) | P(1\|Base) | A | C | G | T |
| --- | --- | --- | --- | --- | --- | --- | --- | --- |
| 240 | C | C | 0.3975 | 0.2358 | 4 | **6492** | 5 | 1 |
| 3036 | C | C | 0.9998 | 0.0001 | 1 | **1422** | 1 | 9 |
| 8781 | C | T | 0.9929 | 0.0012 | 0 | 0 | 0 | **140** |
| 14407 | C | T | 0.0092 | 0.3297 | 1 | 3 | 2 | **7359** |
| 17125 | T | C | 0.0217 | 0.9783 | 1 | **7991** | 0 | 6 |
| 20267 | A | G | 0.0061 | 0.9935 | 0 | 0 | **13** | 0 |
| 23402 | A | G | 0.0066 | 0.3311 | 0 | 0 | **196** | 0 |
| 28143 | T | C | 0.9988 | 0.0005 | 2 | **8004** | 0 | 4 |

Lastly, even though most anomalous sequences reported by Bolotie had clean separations between two parental clades, some mutational signatures contained admixtures in conditional probabilities from other clades. In one such example, illustrated in Figure 2C, at position 240 the conditional probability has a higher affinity towards a blue clade than others. A signature like that could be indicative of a random mutation, sequencing or assembly artifacts, and Bolotie resolved the parental clades in a seemingly parsimonious way.

### Sequencing and assembly artifacts

Although recombination has been extensively observed within and between members of the coronaviridae family, such observations were in the past characterized based on years of accumulated variants in a relatively small collection of genomes. The bulk of the SARS-CoV-2 sequences currently deposited in GISAID and GENBANK were collected and sequenced between the months of March and May of 2020 and show very high degree of similarity, which combined with the large number of available genomes, makes evolutionary analysis very challenging.

Another complication is that an apparent recombinant strain might instead be the result of a co-infection. Suppose a sample was collected from a patient co-infected with two distinct lineages of the virus, where one lineage contains two SNVs while the other lineage contains two distinct SNVs at the same positions. Such a sample would be amplified and sequenced in a single batch. Reads representing both alleles would be provided to an assembler to produce the final genome. Depending on the software, protocols and coverage of each allele by the reads, it is possible that an assembler would produce a sequence with one allele from the first lineage and the other allele from the second lineage, creating an artifactual recombinant. Re-analysis of raw sequencing reads by mapping them against the assembled genome should reveal such artifacts, because reads from a patient co-infected by multiple clades would reveal different variants at the corresponding sites.

To evaluate this hypothesis, we obtained sets of raw sequencing reads deposited at NCBI/SRA or ENA for some of the recombinant genomes identified with Bolotie. Unfortunately, GISAID does not require authors to submit raw data, and only a limited number of submitters have placed their data in public archives such as SRA and ENA. Thus, we were only able to recover a limited number of datasets for our analysis.

One of the sequences for which we were able to identify raw reads was isolate EPI_ISL_439137 (Figure 2B), for which reads were deposited in the European Bioinformatics Institute’s ENA database. As summarized in Table 1, all positions with high conditional affinities for a clade had a homogenous composition, indicating that the data did not derive from two distinct isolates, but instead it was likely a single isolate containing variants from two parental lineages.

Searching the list of recombinant genomes for similar signatures, we identified another isolate, EPI_ISL_489588 also from Scotland, dated one week earlier, which contained the same variants. Another isolate, EPI_ISL_510303 from Spain, had all but one variant (at position 28,143) matching the recombinant signature of the isolates from Scotland. Given the rapid mutation rate in RNA viruses, it is possible that an independent mutation occurred at that position, or that the reference allele is an assembly artifact, however we could not find raw data corresponding to the Spanish isolate and were unable to further investigate possible reasons for the missing variant.

### Performance

Complete analysis of the 87,695 genomes using Bolotie including alignment and index construction took a total of ~5.5 hours using 36 threads on a two 10 core Intel Xeon E5-2680 v2 processors. Using the conditional probability table provided with the software, analyzing a single additional genome takes on average only ~30 seconds.

## Discussion

The method and experiments presented in this work demonstrate that recombination has occurred between the four existing major clades of SARS-CoV-2. While some of the inferred events are likely homoplasies, or could be attributed to technical artifacts, our analysis shows that at least some of the genomes likely represent true cases of recombination.

Of the 225 recombination events identified in our analysis, the majority were represented by single isolates, suggesting that the event was not established in the population. However, because two-thirds of the available genomes were sequenced between late March and early May, it is possible that more data will reveal additional recombinant lineages.

The 225 inferred recombinant genomes comprise less than 1% of all sequences analyzed. It is possible that many more anomalous genomes could be detected by lowering the variant frequency threshold. For example, if we require 50 sequences to confirm a variant rather than 100, the number of informative sites increases more than two-fold from 411 to 996, possibly allowing detection of events that are rarer, such as those which involve smaller emerging lineages within the 4 clades. However, due to decreased specificity such an approach might require stricter manual inspection as the false positive rate is expected to increase substantially.

While overall all events detected by Bolotie passed manual verification, a possibility of mis-assembly in cases of co-infection by particles from different clades could also explain the presence of multiple clade-specific variants within a single genome. Although we were unable to obtain raw read data for all recombinants, our analysis of the EPI_ISL_439137 isolate (Figure 2B, Table 1) shows that at least in one case the recombinant origins of the genome can be validated. Several other recombinants identified in our analysis (EPI_ISL_468407, EPI_ISL_452334, EPI_ISL_475584, EPI_ISL_464547) have also been previously identified by other groups (VanInsberghe et al., 2020).

Due to the differences in library preparation, sequencing technology and assembly protocols, the need for raw data and independent validation is very high. We urge researchers to submit raw sequencing data so that any future studies are able to verify their findings, not only when studying recombination events, but also individual rare variants, transmission patterns, clade prevalence in different populations, etc.

Because our method relies heavily on the accuracy of variant calls, we sought to compare how well available phylogenetic trees agree with trees built using consensus sequences constructed from the alignment data we obtained. The trees shown in figures 3A and 3B are very similar, confirming that consensus sequences constructed by Bolotie preserve essential information sufficient for accurate phylogenetic analysis. In our analysis, Bolotie identified 15 out of 4,039 sequences in the set of sequences used by NextStrain (Hadfield et al., 2018) as recombinants or as having mutational anomalies (Supplementary Table 2). Minor differences between the NextStrain tree and the tree computed from Bolotie consensus sequences are to be expected since consensus sequences have 200 bases replaced with the reference at both 3’ and 5’ ends and do not include any structural variations. Additionally, since NextStrain tree includes 455 isolates submitted to GISAID after we downloaded our latest set, those additional sequences are also expected to slightly alter the topology. Lastly, differences in software versions and randomized methods inherent in the tree-building software are expected to produce trees with minor differences on each iteration.

**Figure 3.**
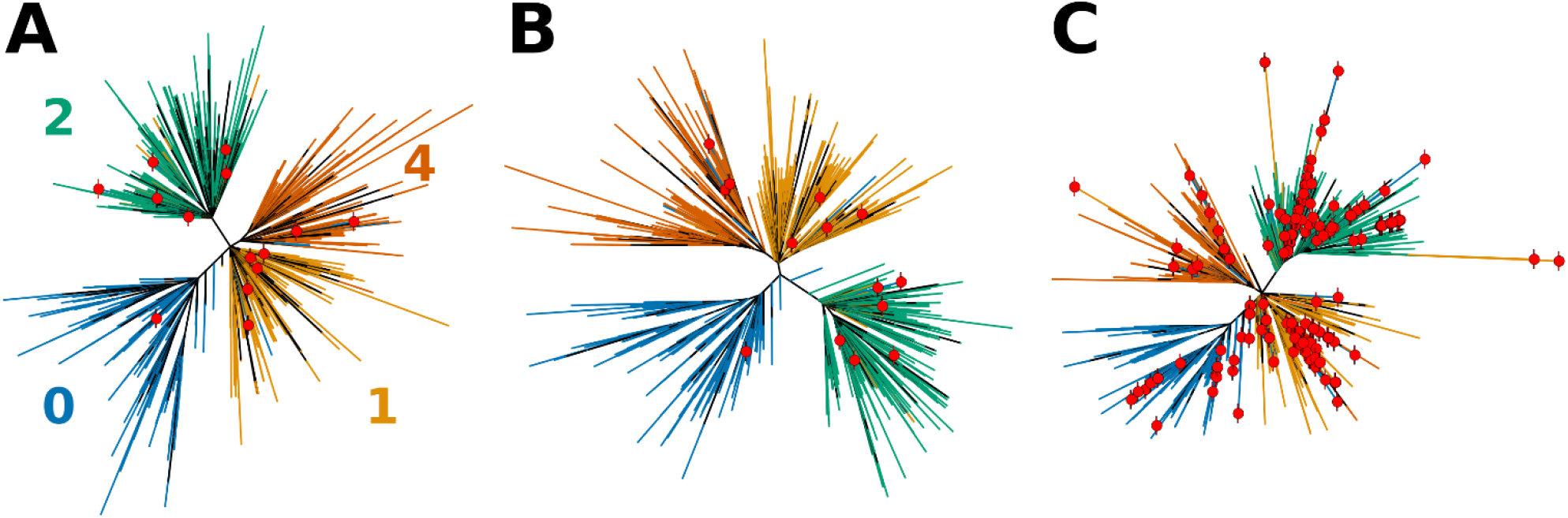
Effects of sequence composition on the topology of the phylogenetic tress for SARS-CoV-2. A tree obtained directly from NextStrain (**A**) is first compared to (**B**) the tree computed using Bolotie consensus sequences for the same set of isolates. **(C)** Shows a tree computed for the same set of isolates with 210 additional recombinant sequences as identified by Bolotie. Leaf nodes that correspond to recombinant genomes are labeled with red dots.

On the other hand, the introduction of 225 recombinant or otherwise anomalous genomes produced a mildly distorted tree with multiple outliers, shown in Figure 3C and Figure 1. In both illustrations most of the recombinant genomes were assigned a clade different from the non-recombinant neighbors by GISAID. Furthermore, several groups of potential recombinant genomes with identical or highly similar mutational signatures were identified by Bolotie (Figure 1). Such lineages are especially important for phylogenetic analysis, as they may affect the topology of the trees more significantly. Even minor perturbations to the topology of the tree in the presence of misclassified outliers may have adverse effects on the studies of dynamics and transmission of the pathogen lineages in the population (Awadalla, 2003). This once again illustrates the importance of properly handling anomalous sequences in phylogenetic analysis.

It must also be noted that different clade assignments for the SARS-CoV-2 genomes currently exist (Hadfield et al., 2018; Shu & McCauley, 2017), at least in part due to differences in tree-building strategies (Supplementary Table 1). However, even though discrepancies in clade assignment may present a challenge to Bolotie, the results will still be of use for the refinement of phylogenetic trees and ultimately clade assignments of the genomes. Grouping together conflicting clades and smaller clades should increase the specificity of the results.

To the best of our knowledge, there currently exists one other recent evaluation of recombination in SARS-CoV-2 genomes (VanInsberghe et al., 2020). Several studies hinted at the possibility of recombination occurring, but the results were inconclusive due to very small sample size at the time (Yi, 2020). Bolotie adds 221 candidate recombinants to the 5 proposed by VanInsberghe et al. Upon comparison of our results we immediately noted the absence of isolate EPI_ISL_464547, one of the 5 genomes reported by VanInsberghe et al., in the output of Bolotie. Of the 5 genomes reported in that study, EPI_ISL_464547 had the shortest length of the second segment. Upon closer investigation of conditional probabilities computed by Bolotie for that genome, we found that none of the variants are clade-defining, nor does the dominant clade 0 ever drop in probability below the baseline of 0.25 at which all genomes are equally likely. The only exception was a variant at the tail end of the sequence, which was equally likely for clades 0, 1 and 3 at 0.33 probability (Supplementary Figure 1). Thus, we concluded that the assignment done by Bolotie was likely correct for that genome.

However, it is possible that differences in partitioning of clades 0, 1 and 3 between our method and that used by VanInsberghe et al. would result in different conditional probabilities at that variant position. As shown in Supplementary Table 1, one such discrepancy does exist between clades 1 and 3, where NextStrain classification adds 334 sequences from clade GH to clade G. However, clade 0 would still have a conditional probability at that position greater than the baseline. As a result, we find it difficult to provide a conclusive assessment of the EPI_ISL_464547 isolate given currently available data.

It must also be stressed that our work focused primarily on the development of a highly efficient, scalable general-purpose method for detecting recombination events in viral genomes irrespective of the divergence rates in the pool of collected isolates. An additional purpose was to search for convincing evidence that recombination does indeed happen in SARS-CoV-2. Hence, we imposed multiple conservative criteria, such as assessment of only 4 clades. However, it is important to note that recombination likely happens within lineages of the same clade. While not targeted in our analysis, future studies may choose to evaluate these intra-clade events, possibly yielding much higher numbers of recombination events.

## Conclusion

Given how recently this novel strain of coronavirus appeared, much remains to be learned about SARS-CoV-2 and how it may change over time. Recombination events, which may have been responsible for the initial emergence of SARS-CoV-2 (Zhang et al., 2020), may have significant impact on future transmission and virulence of the virus. As such, our ability to detect recombination events in a timely manner is crucial in the ongoing efforts to find a solution to the pandemic and prevent additional casualties.

Utilizing an enormous collection of SARS-CoV-2 isolates sequenced by thousands of researchers around the globe (Shu & McCauley, 2017), we were able to develop a method that can reliably detect sequences with anomalous mutation patterns which are indicative of recombination events. Using the proposed method, we identified 225 high likelihood recombinant sequences. Our findings suggest that recombination in SARS-CoV-2 is much more common than previously reported and that several recombinant lineages may have become established in the population.

We hope that the software presented here along with provided pre-built indices will help to detect future recombination events quickly and reliably, and aid in efforts to track the spread of the SARS-CoV-2 virus.

## Methods

In this work we present Bolotie, a collection of methods that enables rapid alignment, variant calling, inter-clade recombination detection, and parent sequence search for large sets of assembled viral genomes. Our method is robust even when the divergence of collected genomes is very small, and it is designed for ultra-fast detection of anomalies. Because Bolotie utilizes a probability index that can be used for a one-against-all analysis, alignments and indices do not have to be recomputed when evaluating novel sequences as recombinants, which greatly increases the efficiency of single-sequence analysis. Our method is designed specifically to facilitate analysis of large datasets of low-divergence sequences, borrowing ideas from recent developments in multiple sequence alignment, phylogenetic analysis and sequence clustering.

### Data

87,695 complete high-coverage genomes were obtained from GISAID (Shu & McCauley, 2017) using the provided interface. SARS-CoV-2 reference genome isolate Wuhan-Hu-1 (GenBank, accession no. <u>MN908947</u>) was obtained from NCBI and used to guide the alignment, variant calling and consensus sequence generation in the protocol.

Clade assignments for the 87,695 available genomes were also obtained from GISAID (Shu & McCauley, 2017). Lineages S, L, V and O were grouped together, similarly to NextStrain (Hadfield et al., 2018), and based on the close distances observed in our independent phylogenetic analyses (Figure 3). Mappings between clade IDs in our analysis, those defined by GISAID, and those defined by NextStrain are provided in Supplementary Table 1.

To test genomes for the possibility that the putative recombinant isolates might represent a co-infection with two or more distinct SARS-CoV-2 isolates, we searched the SRA database of raw read data using all unique GISAID identifiers as well as lab-assigned identifiers extracted from sequence headers.

### Bolotie - Computing the Alignment

A global alignment method to the reference using the implementation from the KSW2 library (Li, 2018; Suzuki & Kasahara, 2018) was developed to facilitate efficient and parallelized alignment and variant calling for each query sequence. Because sequence divergence is very low in the collection of SARS-CoV-2 genomes, the reference-guided approach used instead of the conventional multiple sequence alignment is 1) significantly faster, 2) allows simple addition of new sequences, and 3) does not cause explosions in gaps at 3’ and 5’ ends of the viral genomes. Alignment was performed with a DNAFULL scoring matrix, a gap penalty of 12 and gap extension penalty of 4.

### Bolotie – Computing Consensus Sequences

After the pairwise alignment step, we construct for every genome in the input set a consensus sequence by substituting reference genome alleles for the high-frequency variants called by the aligner. Not only does this approach allow us to filter variants used to search for recombinants, but also produces a set of genomes with a standardized set of coordinates which can be used as a multiple sequence alignment (MSA) for phylogenetic analysis.

Because the 3’ and 5’ ends of viral genomes are notoriously difficult to assemble correctly, we chose to force the first and last 200 bases in the consensus sequence of each genome to be identical to the reference genome in accordance with previous studies (VanInsberghe et al., 2020), regardless of whether SNVs were revealed by alignment in those regions or not.

Next, since Bolotie is designed to work for genomes with very few variants it is particularly sensitive to ambiguous nucleotides. To avoid biases caused by uncalled bases, any such instances were treated as an unknown base (N). Furthermore, to avoid bias in our predictions for all sequences in the dataset we replace nucleotides at a position of low-frequency variants with the reference allele making such positions equally probable for any clade (clade neutral). We define a low-frequency SNV as one that has fewer than 100 genome sequences that differ from the reference sequence at that position.

Because we did not use structural variants when constructing consensus sequences, the final collection of filtered genomes represents a multiple sequence alignment (MSA). This not only allows us to use it directly for phylogenetic analysis but also allow us in the next steps to easily count SNVs across sequences and clades or compute distances among sequences efficiently.

### Bolotie – Identifying Recombinants

Based on the provided clade information for each sequence, a model is created to evaluate each sequence as a potential recombinant. From the MSA the conditional probability for each nucleotide position is computed; i.e., the probability of this position belonging to clade *C*_*i*_ given the nucleotide b. However, to account for differences in clade sizes, we multiply the base counts at each position for each clade with the reciprocal of the number of sequences in that clade. The algorithm also ensures that any ambiguous character is assigned a neutral conditional probability of 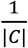. For every position in the MSA we now have the normalized conditional probabilities of that base position belonging to a certain clade given the base observed in the consensus sequence.

Let C = {c_1_, c_2_, …, c_k_} be the clades (in this paper: k=4) with c_i_ ∈ C. We define two sets of bases B_4_={A,C,G,T} and B_16_ being all IUPAC characters. n_ci,b_ denotes the number of sequences in clade c_i_ that have base b ∈ B_4_ at the position of interest. Now the weighted conditional probability Pr(c_i_ | b) of the sequence belonging to clade c_i_ given the observed base b ∈ B_16_ at a certain position is defined as:

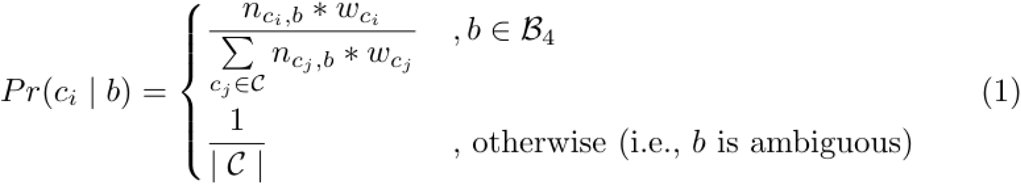

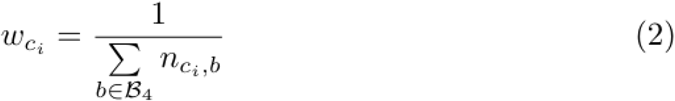

Now we determine for each sequence whether it might be a recombinant. Therefore, we assign each base the most likely clade. To avoid frequent switching of clades or being prone to insignificant differences in probabilities in a single nucleotide, switching between clades from one nucleotide to another is strongly penalized (realized with a transition probability of 0.0001). By multiplying the probabilities along a path, the path with the highest probability can be determined. This can be modeled as an HMM and solved with the Viterbi algorithm in O(nc^2^) where n is the sequence length and c the number of clusters.

At each position of the MSA, each state is representing one clade with the conditional probabilities for every four bases. The Viterbi algorithm then finds the path with the highest likelihood given the sequence of an isolate. The model is identical for every isolate and only depends on the MSA. Figure 4 shows the approach, but only illustrates the conditional probability of the observed base of the sequence in each state.

**Figure 4.**
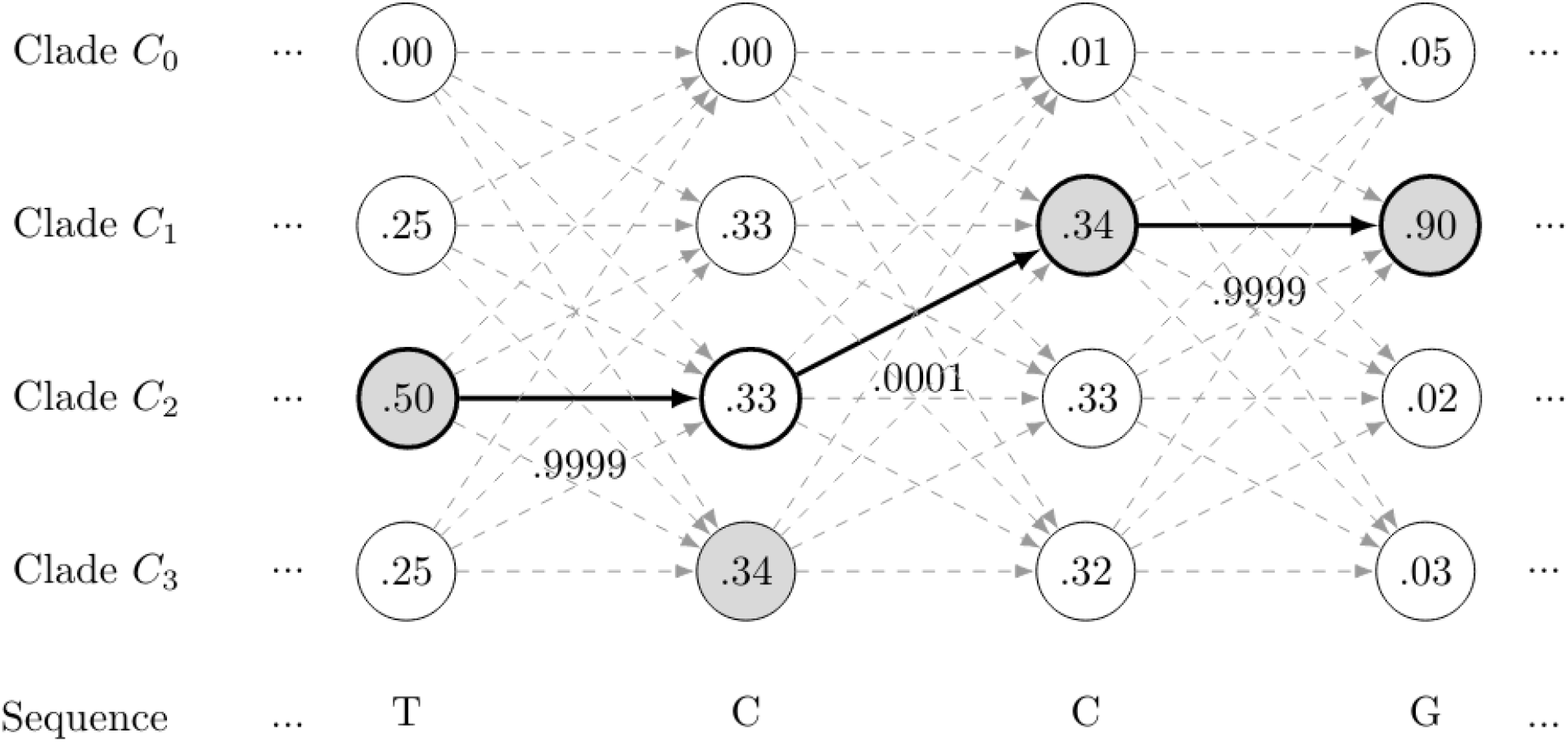
The maximum conditional probability for each nucleotide is highlighted in gray, while the path with the maximum likelihood is highlighted in bold. By penalizing switching of clades, insignificant differences in probabilities between clades as well as short windows representing a switch to a different clade are avoided. For clarity transitions between nodes on non-optimal paths are indicated in gray without labeled probabilities.

This method cannot detect the exact breakpoint location for a recombinant, because it relies on discrete SNV differences between clades; thus, it can only narrow down the breakpoint to the region between two clade-specific SNVs.

### Bolotie – Searching for Closest Parental Genomes

Previous methods for recombination detection examine triplets of sequences to detect a potential recombinant genome and its parents. Unlike such methods, our algorithm relies on conditional probabilities computed for clades – a strategy that provides additional statistical power and also reduces the complexity of the problem.

To identify potential parents of recombinant sequences we have implemented a dedicated method. For each recombined segment of the sequence, as inferred by our algorithm, we compute a Kimura distance matrix (Kimura, 1980) to other sequences in the clade of the corresponding segment. Kimura distance is a distance metric that scores transitions (A <-> G and C <-> T) differently than transversions (interchange of purine for pyrimidine bases). To be more “accurate” we also include low-frequency SNVs that we neglected in the previous step in the consensus sequences. All sequences with the lowest distance score are reported as most likely parents that have contributed the segment in the recombination event.

### Bolotie – Source Code and Data Availability

The core method is implemented in C++ and based on the SeqAn (Reinert et al., 2017) and KSW2 (Li, 2018) libraries, the tree building is performed by IQ-TREE (Minh et al., 2020). The code and test data are available for download on GitHub: github.com/salzberg-lab/bolotie. The SARS-CoV-2 index built using the genomes in our analyses is also available for download at ftp://ftp.ccb.jhu.edu/pub/data/bolotie_sars_cov_2/.

A wrapper script is provided in the GitHub repository to run all steps of the protocol. This script, while convenient, is intended for replicability and testing and lacks some of the available features of Bolotie.

### Bolotie – Parameter Tuning

Our algorithm for path finding is closely inspired by the Viterbi algorithm to search for the most likely path through a sequence of states in a Hidden Markov Model (HMM). Similarly, the algorithm requires setting a probability of transitioning between states, which are clades in our case. While typically estimating this transition probability would be done during training, the scarce accounts of recombination in SARS-CoV-2 were insufficient for such training (VanInsberghe et al., 2020; Yi, 2020). To our knowledge only one study considered a sufficiently large set of genomes, reporting 5 potential recombination events (VanInsberghe et al., 2020). For that reason, instead of determining a transition probability by training a model, we chose to manually inspect the results of several thresholds and manually tune the parameters.

Ideally, however, once enough data is accumulated and more events are verified, a sufficient training set may be available, and a threshold could be determined automatically. Our method is also applicable to the studies of other organisms, such as influenza, where data is more prevalent, and parameters can be tuned with more precision.

### Investigating sequencing data

To test our recombination candidates for signs of co-infection, we aligned available reads with Bowtie2 (Langmead & Salzberg, 2012) against the Wuhan-Hu-1 reference genome using the “--very-sensitive-local" option and otherwise default parameters. The mappings were further sorted and indexed using samtools (Li et al., 2009). Counts for individual nucleotides were obtained using bam-readcount software (https://github.com/genome/bam-readcount) and positions with high conditional probabilities were extracted and summarized.

### Phylogenetic analysis

To test how well information is preserved in our consensus sequences, we obtained a pre-computed tree from NextStrain (Hadfield et al., 2018) which contained a total of 4,494 representative genomes chosen by NextStrain. Only 4,039 genomes from those in the NextStrain tree were available on GISAID at the time when we obtained genome assemblies for our analysis.

First, we re-built the tree using the set of 4,039 genomes using the general time-reversible model as used by the NextStrain platform and allowing IQ-TREE (Minh et al., 2020) to automatically choose the precise model. The same approach was taken to build the phylogenetic tree which included all identified anomalous sequences.

### Visualizations

Visualizations of phylogenetic trees were produced using custom scripts implemented in Python and R. Cladograms illustrated in Figure 3 were constructed in R using the “ape” package (Paradis et al., 2004). Topological circular tree in Figure 1 was produced in Python using the “ETE3” package (Huerta-Cepas et al., 2016) with a custom patch implemented for drawing arcs between leaf nodes.

## Acknowledgements

We would like to thank Dr. Martin Steinegger for the helpful discussions on algorithms for clustering large collection of viral genomes.

