## Supplemental Materials for "Rapid detection of inter-clade recombination in SARS-CoV-2 with Bolotie"

### Supplementary Materials

#### Figures

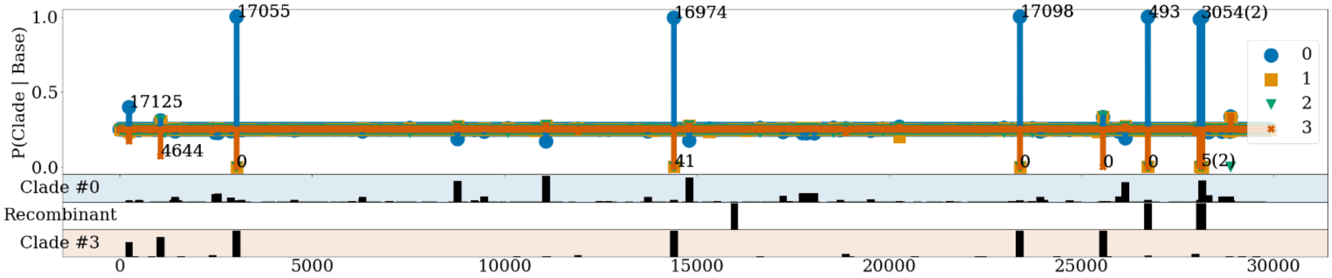


***Supplementary Figure 1.*** *Conditional probability signature of the EPI_ISL_454983 isolate which was not reported as anomalous. The top section shows conditional probabilities of a clade given a nucleotide at each position. Bars are plotted for the two parent clades as reported in* VanInsberghe et al. (VanInsberghe et al., 2020) *and the other clades are shown in dots of corresponding color. Each peak over 0.1 above the baseline (0.25) is labeled with the number of genomes it appears in. An average is reported for multiple variants in close proximity on the plot, listing the number of averaged variants in parentheses. Three bottom sections show the frequency of variants at each position for parental clades (top and bottom rows) and variants observed on the recombinant genome (middle row)*

#### Tables

| **Bolotie** | | **GISAID** | | **NextStrain** | | | | |
|---|---|---|---|---|---|---|---|---|
| ***Cluster*** | ***# Genomes*** | ***Clade*** | ***# Genomes*** | ***19A*** | ***19B*** | ***20A*** | ***20B*** | ***20C*** |
| 0 | 1059 | L | 199 | 199 | 0 | 0 | 0 | 0 |
| | | S | 279 | 3 | 276 | 0 | 0 | 0 |
| | | V | 113 | 113 | 0 | 0 | 0 | 0 |
| | | O | 468 | 394 | 13 | 36 | 23 | 2 |
| 1 | 1005 | G | 1005 | 9 | 0 | 939 | 50 | 7 |
| 2 | 1203 | GR | 1203 | 0 | 0 | 0 | 1203 | 0 |
| 3 | 772 | GH | 772 | 0 | 0 | 334 | 1 | 437 |

***Supplementary Table 1.*** *Mappings between clade identifiers in our analysis and those defined by GISAID* (Shu & McCauley, 2017) *and NextStrain* (Hadfield et al., 2018). *Only 4,494 sequences for which NextStrain assignments were available were counted for this table. Clades in our analysis were taken directly from GISAID, however, smaller lineages L,S,V as well as sequences labeled as "other" (O) were grouped together as sharing neighborhood on the tree.*

| **Sequence Identifier** |
|---|
| hCoV-19_Turkey_KOU-MG_2020_EPI_ISL_476832_2020-04-27 |
| hCoV-19_Bahrain_920268866_2020_EPI_ISL_510531_2020-06-23 |
| hCoV-19_SaudiArabia_KAUST-JEDDAH888_2020_EPI_ISL_513037_2020-05-24 |
| hCoV-19_Uganda_UG003_2020_EPI_ISL_451185_2020-03-27 |
| hCoV-19_Bosnia_and_Herzegovina_01-Livno_2020_EPI_ISL_462753_2020-05-27 |
| hCoV-19_Russia_StPetersburg-RII8935S_2020_EPI_ISL_450292_2020-04-21 |
| hCoV-19_Austria_Graz-MUG9_2020_EPI_ISL_437299_2020-04-09 |
| hCoV-19_Austria_Graz-MUG3_2020_EPI_ISL_437199_2020-03-31 |
| hCoV-19_Croatia_297_Varazdin_2020_EPI_ISL_454592_2020-03-05 |

| |
|---|
| hCoV-19_Mali_M002673_2020_EPI_ISL_487455_2020-04-10 |
| hCoV-19_Croatia_1761_Dubrovnik_2020_EPI_ISL_454588_2020-03-20 |
| hCoV-19_Czech_Republic_IAB_1_2020_EPI_ISL_426883_2020-03-27 |
| hCoV-19_Oman_RESP-20-4943_2020_EPI_ISL_525425_2020-03-23 |
| hCoV-19_Gambia_0214_2020_EPI_ISL_471158_2020-03-29 |
| hCoV-19_SaudiArabia_KAUST-MADINAH609_2020_EPI_ISL_513172_2020-05-06 |

***Supplementary Table 2.*** *List of SARS-CoV-2 genomes flagged by Bolotie as recombinant out of 4,494 representative isolates present in the NextStrain tree at the time of the analysis.*

| Clade | # Recombinants | # Parents |
|---|---|---|
| 0 | 109 | 171 |
| 1 | 111 | 41 |
| 2 | 5 | 148 |
| 3 | 0 | 90 |
| Total | 225 | 450 |

***Supplementary Table 3.*** *Summary of the GISAID clade assignment of recombinant genomes and inferred parents.*

| **Recombinant** | **Parent #1** | | | **Parent #2** | | |
|---|---|---|---|---|---|---|
| ***Recombinant*** | ***Clade*** | ***Segment*** | ***GISAID*** | ***Clade*** | ***Segment*** | ***GISAID*** |
| EPI_ISL_453713 | 0 | 0-4389 | EPI_ISL_474909 | 2 | 4389-29902 | EPI_ISL_457724 |
| EPI_ISL_453712 | 0 | 0-4389 | EPI_ISL_522572 | 1 | 4389-29902 | EPI_ISL_442358 |
| EPI_ISL_481145 | 0 | 0-4389 | EPI_ISL_422772 | 1 | 4389-29902 | EPI_ISL_458053 |
| EPI_ISL_478027 | 0 | 0-14804 | EPI_ISL_459691 | 1 | 14804-29902 | EPI_ISL_477884 |
| EPI_ISL_425846 | 0 | 0-4389 | EPI_ISL_414593 | 1 | 4389-29902 | EPI_ISL_469073 |
| EPI_ISL_453753 | 0 | 0-4389 | EPI_ISL_474909 | 2 | 4389-29902 | EPI_ISL_420261 |
| EPI_ISL_453752 | 0 | 0-4389 | EPI_ISL_417203 | 1 | 4389-29902 | EPI_ISL_442479 |
| EPI_ISL_489588 | 0 | 0-8916 | EPI_ISL_488600 | 1 | 8916-29902 | EPI_ISL_428679 |
| EPI_ISL_453714 | 0 | 0-4389 | EPI_ISL_522572 | 2 | 4389-29902 | EPI_ISL_418317 |
| EPI_ISL_510531 | 0 | 0-4389 | EPI_ISL_500635 | 2 | 4389-29902 | EPI_ISL_496559 |
| EPI_ISL_479572 | 0 | 0-28656 | EPI_ISL_447565 | 2 | 28656-29902 | EPI_ISL_465904 |
| EPI_ISL_453729 | 0 | 0-4389 | EPI_ISL_417203 | 2 | 4389-29902 | EPI_ISL_462249 |
| EPI_ISL_491989 | 0 | 0-4389 | EPI_ISL_517362 | 2 | 4389-29902 | EPI_ISL_435124 |
| EPI_ISL_507986 | 3 | 0-27877 | EPI_ISL_428790 | 2 | 27877-29902 | EPI_ISL_421199 |
| EPI_ISL_422850 | 1 | 0-20134 | EPI_ISL_510174 | 0 | 20134-29902 | EPI_ISL_455008 |
| EPI_ISL_437472 | 1 | 0-20134 | EPI_ISL_428687 | 0 | 20134-29902 | EPI_ISL_433299 |
| EPI_ISL_513037 | 3 | 0-27877 | EPI_ISL_436478 | 2 | 27877-29902 | EPI_ISL_443672 |
| EPI_ISL_459371 | 3 | 0-27877 | EPI_ISL_417336 | 2 | 27877-29902 | EPI_ISL_443781 |
| EPI_ISL_450292 | 1 | 0-14877 | EPI_ISL_423026 | 0 | 14877-29902 | EPI_ISL_462185 |
| EPI_ISL_496321 | 2 | 0-20980 | EPI_ISL_495677 | 1 | 20980-29902 | EPI_ISL_443697 |
| EPI_ISL_497594 | 2 | 0-20980 | EPI_ISL_497589 | 1 | 20980-29902 | EPI_ISL_439416 |
| EPI_ISL_497455 | 2 | 0-20980 | EPI_ISL_496222 | 1 | 20980-29902 | EPI_ISL_447718 |
| EPI_ISL_496334 | 2 | 0-21614 | EPI_ISL_495706 | 1 | 21614-29902 | EPI_ISL_426986 |
| EPI_ISL_525424 | 2 | 0-18486 | EPI_ISL_491128 | 0 | 18486-29902 | EPI_ISL_416044 |
| EPI_ISL_497021 | 0 | 0-14804 | EPI_ISL_437300 | 2 | 14804-29902 | EPI_ISL_423582 |
| EPI_ISL_525428 | 2 | 0-18486 | EPI_ISL_458116 | 0 | 18486-29902 | EPI_ISL_416580 |
| EPI_ISL_468407 | 3 | 0-18736 | EPI_ISL_418037 | 0 | 18736-29902 | EPI_ISL_426512 |
| EPI_ISL_508098 | 3 | 0-27384 | EPI_ISL_429316 | 2 | 27384-29902 | EPI_ISL_517083 |
| EPI_ISL_520686 | 1 | 0-20134 | EPI_ISL_457985 | 0 | 20134-29902 | EPI_ISL_483126 |

| | | | | | | |
|---|---|---|---|---|---|---|
| EPI_ISL_495767 | 0 | 0-28310 | EPI_ISL_463692 | 2 | 28310-29902 | EPI_ISL_481807 |
| EPI_ISL_476832 | 0 | 0-14804 | EPI_ISL_426680 | 1 | 14804-29902 | EPI_ISL_417790 |
| EPI_ISL_450122 | 0 | 0-4389 | EPI_ISL_454919 | 1 | 4389-29902 | EPI_ISL_460086 |
| EPI_ISL_420268 | 0 | 0-4389 | EPI_ISL_436738 | 2 | 4389-29902 | EPI_ISL_420278 |
| EPI_ISL_438422 | 3 | 0-27877 | EPI_ISL_418425 | 2 | 27877-29902 | EPI_ISL_436293 |
| EPI_ISL_518315 | 2 | 0-28878 | EPI_ISL_520048 | 0 | 28878-29902 | EPI_ISL_479960 |
| EPI_ISL_487348 | 2 | 0-28869 | EPI_ISL_482867 | 0 | 28869-29902 | EPI_ISL_427453 |
| EPI_ISL_518193 | 2 | 0-28878 | EPI_ISL_522218 | 0 | 28878-29902 | EPI_ISL_518855 |
| EPI_ISL_508009 | 3 | 0-27877 | EPI_ISL_508873 | 2 | 27877-29902 | EPI_ISL_523989 |
| EPI_ISL_520527 | 2 | 0-28878 | EPI_ISL_521787 | 0 | 28878-29902 | EPI_ISL_499667 |
| EPI_ISL_519690 | 2 | 0-28878 | EPI_ISL_521589 | 0 | 28878-29902 | EPI_ISL_479960 |
| EPI_ISL_504177 | 3 | 0-27877 | EPI_ISL_513259 | 2 | 27877-29902 | EPI_ISL_469830 |
| EPI_ISL_519112 | 2 | 0-28878 | EPI_ISL_521787 | 0 | 28878-29902 | EPI_ISL_499667 |
| EPI_ISL_521605 | 2 | 0-28878 | EPI_ISL_522114 | 0 | 28878-29902 | EPI_ISL_479960 |
| EPI_ISL_512969 | 3 | 0-27877 | EPI_ISL_512953 | 2 | 27877-29902 | EPI_ISL_480264 |
| EPI_ISL_475026 | 3 | 0-27877 | EPI_ISL_418430 | 2 | 27877-29902 | EPI_ISL_454064 |
| EPI_ISL_518953 | 2 | 0-28878 | EPI_ISL_521585 | 0 | 28878-29902 | EPI_ISL_499667 |
| EPI_ISL_521775 | 2 | 0-28878 | EPI_ISL_521796 | 0 | 28878-29902 | EPI_ISL_479960 |
| EPI_ISL_512983 | 0 | 0-3095 | EPI_ISL_437716 | 3 | 3095-29902 | EPI_ISL_512981 |
| EPI_ISL_436055 | 0 | 0-3095 | EPI_ISL_482575 | 3 | 3095-29902 | EPI_ISL_420797 |
| EPI_ISL_521286 | 2 | 0-28878 | EPI_ISL_518623 | 0 | 28878-29902 | EPI_ISL_471263 |
| EPI_ISL_464140 | 0 | 0-23928 | EPI_ISL_421655 | 2 | 23928-29902 | EPI_ISL_423851 |
| EPI_ISL_519756 | 2 | 0-28878 | EPI_ISL_519506 | 0 | 28878-29902 | EPI_ISL_438065 |
| EPI_ISL_521351 | 2 | 0-28878 | EPI_ISL_519476 | 0 | 28878-29902 | EPI_ISL_438065 |
| EPI_ISL_479518 | 1 | 0-24075 | EPI_ISL_461481 | 2 | 24075-29902 | EPI_ISL_459379 |
| EPI_ISL_521825 | 2 | 0-28878 | EPI_ISL_522065 | 0 | 28878-29902 | EPI_ISL_518855 |
| EPI_ISL_518970 | 2 | 0-28878 | EPI_ISL_519303 | 0 | 28878-29902 | EPI_ISL_499667 |
| EPI_ISL_512950 | 0 | 0-3095 | EPI_ISL_437716 | 3 | 3095-29902 | EPI_ISL_437486 |
| EPI_ISL_520655 | 2 | 0-28878 | EPI_ISL_521787 | 0 | 28878-29902 | EPI_ISL_499667 |
| EPI_ISL_516778 | 3 | 0-25563 | EPI_ISL_513293 | 2 | 25563-29902 | EPI_ISL_521074 |
| EPI_ISL_517938 | 3 | 0-27877 | EPI_ISL_514211 | 2 | 27877-29902 | EPI_ISL_425361 |
| EPI_ISL_520605 | 2 | 0-28878 | EPI_ISL_520165 | 0 | 28878-29902 | EPI_ISL_499667 |
| EPI_ISL_496298 | 0 | 0-28310 | EPI_ISL_461482 | 2 | 28310-29902 | EPI_ISL_447710 |
| EPI_ISL_474976 | 3 | 0-27877 | EPI_ISL_436059 | 2 | 27877-29424 | EPI_ISL_500061 |
| EPI_ISL_496254 | 3 | 0-24047 | EPI_ISL_449979 | 0 | 24047-29902 | EPI_ISL_415621 |
| EPI_ISL_470879 | 3 | 0-9724 | EPI_ISL_460103 | 0 | 9724-29902 | EPI_ISL_447561 |
| EPI_ISL_519335 | 2 | 0-28878 | EPI_ISL_521589 | 0 | 28878-29902 | EPI_ISL_479960 |
| EPI_ISL_455354 | 0 | 0-8916 | EPI_ISL_455352 | 1 | 8916-29902 | EPI_ISL_414443 |
| EPI_ISL_512062 | 3 | 0-28878 | EPI_ISL_447033 | 2 | 28878-29902 | EPI_ISL_462369 |
| EPI_ISL_520553 | 2 | 0-28878 | EPI_ISL_521067 | 0 | 28878-29902 | EPI_ISL_471263 |
| EPI_ISL_519186 | 2 | 0-28878 | EPI_ISL_519279 | 0 | 28878-29902 | EPI_ISL_499667 |
| EPI_ISL_516769 | 3 | 0-27877 | EPI_ISL_418036 | 2 | 27877-29902 | EPI_ISL_494082 |
| EPI_ISL_426890 | 3 | 0-27877 | EPI_ISL_417830 | 2 | 27877-29902 | EPI_ISL_426356 |
| EPI_ISL_483142 | 0 | 0-4389 | EPI_ISL_416578 | 1 | 4389-29902 | EPI_ISL_419767 |
| EPI_ISL_513052 | 0 | 0-3095 | EPI_ISL_475666 | 3 | 3095-29902 | EPI_ISL_513184 |
| EPI_ISL_521728 | 2 | 0-28878 | EPI_ISL_521796 | 0 | 28878-29902 | EPI_ISL_479960 |
| EPI_ISL_520644 | 2 | 0-28878 | EPI_ISL_521036 | 0 | 28878-29902 | EPI_ISL_479960 |
| EPI_ISL_495724 | 2 | 0-28881 | EPI_ISL_495647 | 0 | 28881-29902 | EPI_ISL_514665 |
| EPI_ISL_483136 | 3 | 0-27877 | EPI_ISL_498042 | 2 | 27877-29902 | EPI_ISL_472195 |
| EPI_ISL_519785 | 2 | 0-28878 | EPI_ISL_521813 | 0 | 28878-29902 | EPI_ISL_438065 |

| | | | | | | |
|---|---|---|---|---|---|---|
| EPI_ISL_467431 | 2 | 0-13730 | EPI_ISL_475093 | 0 | 13730-29902 | EPI_ISL_419791 |
| EPI_ISL_519970 | 2 | 0-28878 | EPI_ISL_519312 | 0 | 28878-29902 | EPI_ISL_438065 |
| EPI_ISL_521794 | 2 | 0-28878 | EPI_ISL_522240 | 0 | 28878-29902 | EPI_ISL_479960 |
| EPI_ISL_519895 | 2 | 0-28878 | EPI_ISL_522065 | 0 | 28878-29902 | EPI_ISL_499667 |
| EPI_ISL_513211 | 0 | 0-3095 | EPI_ISL_513162 | 3 | 3095-29902 | EPI_ISL_513171 |
| EPI_ISL_517907 | 3 | 0-27877 | EPI_ISL_513293 | 2 | 27877-29902 | EPI_ISL_466545 |
| EPI_ISL_417420 | 3 | 0-20270 | EPI_ISL_420307 | 0 | 20270-29902 | EPI_ISL_416405 |
| EPI_ISL_430339 | 1 | 0-20134 | EPI_ISL_427772 | 0 | 20134-29902 | EPI_ISL_482320 |
| EPI_ISL_475595 | 0 | 0-14804 | EPI_ISL_426680 | 3 | 14804-29902 | EPI_ISL_508883 |
| EPI_ISL_519129 | 2 | 0-28878 | EPI_ISL_520891 | 0 | 28878-29902 | EPI_ISL_499667 |
| EPI_ISL_462753 | 0 | 0-4389 | EPI_ISL_422816 | 2 | 4389-29902 | EPI_ISL_450255 |
| EPI_ISL_512937 | 3 | 0-27877 | EPI_ISL_455778 | 2 | 27877-29902 | EPI_ISL_431909 |
| EPI_ISL_516777 | 3 | 0-25563 | EPI_ISL_513293 | 2 | 25563-29902 | EPI_ISL_521074 |
| EPI_ISL_525425 | 2 | 0-18486 | EPI_ISL_445270 | 0 | 18486-29902 | EPI_ISL_443222 |
| EPI_ISL_454425 | 1 | 0-26144 | EPI_ISL_460084 | 0 | 26144-29902 | EPI_ISL_413855 |
| EPI_ISL_423475 | 3 | 0-27877 | EPI_ISL_418377 | 2 | 27877-29902 | EPI_ISL_438074 |
| EPI_ISL_513039 | 3 | 0-27877 | EPI_ISL_479854 | 2 | 27877-29902 | EPI_ISL_443672 |
| EPI_ISL_513172 | 3 | 0-20270 | EPI_ISL_513148 | 0 | 20270-29902 | EPI_ISL_440724 |
| EPI_ISL_512943 | 0 | 0-3095 | EPI_ISL_437299 | 3 | 3095-29902 | EPI_ISL_512918 |
| EPI_ISL_516767 | 3 | 0-27877 | EPI_ISL_513293 | 2 | 27877-29902 | EPI_ISL_511415 |
| EPI_ISL_459916 | 0 | 0-25562 | EPI_ISL_419761 | 2 | 25562-29902 | EPI_ISL_428920 |
| EPI_ISL_426883 | 3 | 0-27877 | EPI_ISL_525541 | 2 | 27877-29902 | EPI_ISL_513512 |
| EPI_ISL_494085 | 3 | 0-7600 | EPI_ISL_456033 | 2 | 7600-29902 | EPI_ISL_474046 |
| EPI_ISL_514281 | 1 | 0-28880 | EPI_ISL_475682 | 2 | 28880-29902 | EPI_ISL_510722 |
| EPI_ISL_519165 | 2 | 0-28878 | EPI_ISL_519312 | 0 | 28878-29902 | EPI_ISL_438065 |
| EPI_ISL_447591 | 0 | 0-14804 | EPI_ISL_482575 | 2 | 14804-29902 | EPI_ISL_488199 |
| EPI_ISL_513190 | 3 | 0-23185 | EPI_ISL_513163 | 0 | 23185-29902 | EPI_ISL_513033 |
| EPI_ISL_518372 | 2 | 0-28878 | EPI_ISL_519303 | 0 | 28878-29902 | EPI_ISL_499667 |
| EPI_ISL_515358 | 3 | 0-17470 | EPI_ISL_515319 | 0 | 17470-29902 | EPI_ISL_482471 |
| EPI_ISL_521595 | 2 | 0-28878 | EPI_ISL_522225 | 0 | 28878-29902 | EPI_ISL_479960 |
| EPI_ISL_525618 | 0 | 0-14804 | EPI_ISL_418159 | 3 | 14804-29902 | EPI_ISL_429266 |
| EPI_ISL_483857 | 3 | 0-24047 | EPI_ISL_469047 | 0 | 24047-29902 | EPI_ISL_434171 |
| EPI_ISL_509704 | 0 | 0-3095 | EPI_ISL_437299 | 3 | 3095-29902 | EPI_ISL_436635 |
| EPI_ISL_521397 | 2 | 0-28878 | EPI_ISL_518123 | 0 | 28878-29902 | EPI_ISL_499667 |
| EPI_ISL_521620 | 2 | 0-28878 | EPI_ISL_522101 | 0 | 28878-29902 | EPI_ISL_479960 |
| EPI_ISL_446166 | 0 | 0-14723 | EPI_ISL_426680 | 1 | 14723-29902 | EPI_ISL_422018 |
| EPI_ISL_437202 | 0 | 0-3095 | EPI_ISL_421911 | 3 | 3095-29902 | EPI_ISL_475921 |
| EPI_ISL_513000 | 0 | 0-3095 | EPI_ISL_437199 | 3 | 3095-29902 | EPI_ISL_437750 |
| EPI_ISL_512956 | 3 | 0-27877 | EPI_ISL_444276 | 2 | 27877-29902 | EPI_ISL_417829 |
| EPI_ISL_456315 | 3 | 0-18736 | EPI_ISL_428388 | 0 | 18736-29902 | EPI_ISL_417106 |
| EPI_ISL_521623 | 2 | 0-28878 | EPI_ISL_522253 | 0 | 28878-29902 | EPI_ISL_518855 |
| EPI_ISL_447688 | 0 | 0-3095 | EPI_ISL_509695 | 3 | 3095-29902 | EPI_ISL_450035 |
| EPI_ISL_454588 | 0 | 0-4389 | EPI_ISL_450235 | 1 | 4389-29902 | EPI_ISL_454583 |
| EPI_ISL_521682 | 2 | 0-28878 | EPI_ISL_522238 | 0 | 28878-29902 | EPI_ISL_499667 |
| EPI_ISL_520604 | 2 | 0-28878 | EPI_ISL_521836 | 0 | 28878-29902 | EPI_ISL_471263 |
| EPI_ISL_453710 | 0 | 0-4389 | EPI_ISL_455626 | 2 | 4389-29902 | EPI_ISL_453736 |
| EPI_ISL_511512 | 2 | 0-28878 | EPI_ISL_420400 | 0 | 28878-29902 | EPI_ISL_467132 |
| EPI_ISL_487362 | 3 | 0-27877 | EPI_ISL_466638 | 2 | 27877-29902 | EPI_ISL_475167 |
| EPI_ISL_435666 | 0 | 0-3095 | EPI_ISL_455352 | 3 | 3095-29902 | EPI_ISL_424855 |
| EPI_ISL_513021 | 0 | 0-23928 | EPI_ISL_512995 | 2 | 23928-29902 | EPI_ISL_426892 |

| | | | | | | |
|---|---|---|---|---|---|---|
| EPI_ISL_487337 | 1 | 0-28835 | EPI_ISL_467479 | 2 | 28835-29902 | EPI_ISL_420719 |
| EPI_ISL_521598 | 2 | 0-28878 | EPI_ISL_520179 | 0 | 28878-29902 | EPI_ISL_479960 |
| EPI_ISL_451967 | 1 | 0-20134 | EPI_ISL_467148 | 0 | 20134-29902 | EPI_ISL_416353 |
| EPI_ISL_471158 | 3 | 0-27877 | EPI_ISL_476814 | 2 | 27877-29902 | EPI_ISL_436293 |
| EPI_ISL_521759 | 2 | 0-28878 | EPI_ISL_521452 | 0 | 28878-29902 | EPI_ISL_518855 |
| EPI_ISL_445110 | 0 | 0-12477 | EPI_ISL_415743 | 3 | 12477-29902 | EPI_ISL_415481 |
| EPI_ISL_497312 | 2 | 0-27045 | EPI_ISL_463454 | 3 | 27045-29902 | EPI_ISL_524078 |
| EPI_ISL_495049 | 3 | 0-28878 | EPI_ISL_476873 | 2 | 28878-29902 | EPI_ISL_465255 |
| EPI_ISL_481148 | 0 | 0-12477 | EPI_ISL_421911 | 3 | 12477-29902 | EPI_ISL_437452 |
| EPI_ISL_487455 | 1 | 0-20134 | EPI_ISL_420368 | 0 | 20134-29902 | EPI_ISL_417313 |
| EPI_ISL_520595 | 2 | 0-28878 | EPI_ISL_521589 | 0 | 28878-29902 | EPI_ISL_479960 |
| EPI_ISL_519162 | 2 | 0-28878 | EPI_ISL_521777 | 0 | 28878-29902 | EPI_ISL_438065 |
| EPI_ISL_437298 | 0 | 0-3095 | EPI_ISL_509700 | 3 | 3095-29902 | EPI_ISL_475921 |
| EPI_ISL_447679 | 0 | 0-14804 | EPI_ISL_426680 | 2 | 14804-29902 | EPI_ISL_447749 |
| EPI_ISL_513173 | 0 | 0-8916 | EPI_ISL_475584 | 3 | 8916-29902 | EPI_ISL_437486 |
| EPI_ISL_449266 | 0 | 0-4389 | EPI_ISL_499625 | 1 | 4389-29902 | EPI_ISL_433110 |
| EPI_ISL_435662 | 0 | 0-14804 | EPI_ISL_429104 | 3 | 14804-29902 | EPI_ISL_460366 |
| EPI_ISL_493085 | 0 | 0-3095 | EPI_ISL_447568 | 3 | 3095-29902 | EPI_ISL_426556 |
| EPI_ISL_475584 | 0 | 0-23928 | EPI_ISL_429084 | 2 | 23928-29902 | EPI_ISL_416519 |
| EPI_ISL_452334 | 0 | 0-23928 | EPI_ISL_482575 | 2 | 23928-29902 | EPI_ISL_454023 |
| EPI_ISL_455352 | 0 | 0-8916 | EPI_ISL_452761 | 1 | 8916-29902 | EPI_ISL_420551 |
| EPI_ISL_495041 | 3 | 0-27877 | EPI_ISL_476858 | 2 | 27877-29902 | EPI_ISL_465859 |
| EPI_ISL_455773 | 0 | 0-14804 | EPI_ISL_482575 | 3 | 14804-29902 | EPI_ISL_513233 |
| EPI_ISL_521323 | 2 | 0-28878 | EPI_ISL_521731 | 0 | 28878-29902 | EPI_ISL_438065 |
| EPI_ISL_485810 | 1 | 0-20134 | EPI_ISL_478003 | 0 | 20134-29902 | EPI_ISL_439941 |
| EPI_ISL_513012 | 0 | 0-23928 | EPI_ISL_512995 | 2 | 23928-29902 | EPI_ISL_507037 |
| EPI_ISL_514306 | 3 | 0-28851 | EPI_ISL_514284 | 2 | 28851-29424 | EPI_ISL_463452 |
| EPI_ISL_513232 | 0 | 0-8916 | EPI_ISL_475584 | 3 | 8916-29902 | EPI_ISL_437748 |
| EPI_ISL_475589 | 0 | 0-14804 | EPI_ISL_482575 | 3 | 14804-29902 | EPI_ISL_424975 |
| EPI_ISL_457320 | 0 | 0-27383 | EPI_ISL_425845 | 2 | 27383-29902 | EPI_ISL_500101 |
| EPI_ISL_513197 | 0 | 0-3095 | EPI_ISL_513178 | 3 | 3095-29902 | EPI_ISL_430819 |
| EPI_ISL_435073 | 0 | 0-3095 | EPI_ISL_451352 | 3 | 3095-29902 | EPI_ISL_437326 |
| EPI_ISL_512931 | 0 | 0-3095 | EPI_ISL_509700 | 3 | 3095-29902 | EPI_ISL_513233 |
| EPI_ISL_508096 | 1 | 0-18736 | EPI_ISL_510298 | 0 | 18736-29902 | EPI_ISL_463509 |
| EPI_ISL_513182 | 0 | 0-3095 | EPI_ISL_437199 | 3 | 3095-29902 | EPI_ISL_454802 |
| EPI_ISL_519256 | 2 | 0-28878 | EPI_ISL_518473 | 0 | 28878-29902 | EPI_ISL_479960 |
| EPI_ISL_437200 | 0 | 0-3095 | EPI_ISL_509695 | 3 | 3095-29902 | EPI_ISL_475921 |
| EPI_ISL_449271 | 0 | 0-4389 | EPI_ISL_451318 | 1 | 4389-29902 | EPI_ISL_420152 |
| EPI_ISL_516754 | 3 | 0-25433 | EPI_ISL_514177 | 1 | 25433-29902 | EPI_ISL_437751 |
| EPI_ISL_513168 | 0 | 0-3095 | EPI_ISL_437745 | 3 | 3095-29902 | EPI_ISL_444472 |
| EPI_ISL_437745 | 0 | 0-3095 | EPI_ISL_452334 | 3 | 3095-29902 | EPI_ISL_437483 |
| EPI_ISL_454592 | 2 | 0-15438 | EPI_ISL_416140 | 1 | 15438-29902 | EPI_ISL_450198 |
| EPI_ISL_508044 | 3 | 0-27384 | EPI_ISL_512795 | 2 | 27384-29902 | EPI_ISL_511535 |
| EPI_ISL_437203 | 0 | 0-14804 | EPI_ISL_426680 | 1 | 14804-29902 | EPI_ISL_516988 |
| EPI_ISL_449261 | 0 | 0-4389 | EPI_ISL_451513 | 1 | 4389-29902 | EPI_ISL_426009 |
| EPI_ISL_483871 | 3 | 0-27877 | EPI_ISL_476858 | 0 | 27877-29902 | EPI_ISL_428893 |
| EPI_ISL_513046 | 0 | 0-23928 | EPI_ISL_512995 | 2 | 23928-29902 | EPI_ISL_487933 |
| EPI_ISL_463669 | 0 | 0-14804 | EPI_ISL_482575 | 2 | 14804-29902 | EPI_ISL_428895 |
| EPI_ISL_521621 | 2 | 0-28878 | EPI_ISL_520175 | 0 | 28878-29902 | EPI_ISL_518855 |
| EPI_ISL_519626 | 2 | 0-28878 | EPI_ISL_518963 | 0 | 28878-29902 | EPI_ISL_479960 |

| | | | | | | |
|---|---|---|---|---|---|---|
| EPI_ISL_467625 | 0 | 0-8916 | EPI_ISL_493179 | 3 | 8916-29902 | EPI_ISL_458020 |
| EPI_ISL_518935 | 2 | 0-28878 | EPI_ISL_519266 | 0 | 28878-29902 | EPI_ISL_441368 |
| EPI_ISL_519468 | 2 | 0-28878 | EPI_ISL_480641 | 0 | 28878-29902 | EPI_ISL_499667 |
| EPI_ISL_483706 | 3 | 0-11109 | EPI_ISL_444586 | 0 | 11109-29902 | EPI_ISL_416629 |
| EPI_ISL_521572 | 2 | 0-28878 | EPI_ISL_521853 | 0 | 28878-29902 | EPI_ISL_479960 |
| EPI_ISL_516779 | 3 | 0-27877 | EPI_ISL_419544 | 2 | 27877-29902 | EPI_ISL_445315 |
| EPI_ISL_508020 | 3 | 0-27384 | EPI_ISL_508873 | 2 | 27384-29902 | EPI_ISL_480154 |
| EPI_ISL_497082 | 3 | 0-27877 | EPI_ISL_495771 | 2 | 27877-29552 | EPI_ISL_458466 |
| EPI_ISL_420343 | 3 | 0-27877 | EPI_ISL_429968 | 2 | 27877-29902 | EPI_ISL_437998 |
| EPI_ISL_522541 | 3 | 0-24047 | EPI_ISL_510588 | 0 | 24047-29902 | EPI_ISL_427813 |
| EPI_ISL_509874 | 1 | 0-19838 | EPI_ISL_448101 | 2 | 19838-29902 | EPI_ISL_458229 |
| EPI_ISL_521846 | 2 | 0-28878 | EPI_ISL_480641 | 0 | 28878-29902 | EPI_ISL_518855 |
| EPI_ISL_493070 | 1 | 0-20134 | EPI_ISL_439344 | 0 | 20134-29902 | EPI_ISL_422786 |
| EPI_ISL_507011 | 3 | 0-27877 | EPI_ISL_416753 | 2 | 27877-29902 | EPI_ISL_464231 |
| EPI_ISL_519455 | 2 | 0-28878 | EPI_ISL_480641 | 0 | 28878-29902 | EPI_ISL_471263 |
| EPI_ISL_521324 | 2 | 0-28878 | EPI_ISL_520526 | 0 | 28878-29902 | EPI_ISL_438065 |
| EPI_ISL_476267 | 0 | 0-4389 | EPI_ISL_500401 | 2 | 4389-29902 | EPI_ISL_515541 |
| EPI_ISL_449259 | 0 | 0-4389 | EPI_ISL_415609 | 2 | 4389-29902 | EPI_ISL_439243 |
| EPI_ISL_520948 | 2 | 0-28878 | EPI_ISL_520268 | 0 | 28878-29902 | EPI_ISL_479960 |
| EPI_ISL_520810 | 2 | 0-28878 | EPI_ISL_522217 | 0 | 28878-29902 | EPI_ISL_479960 |
| EPI_ISL_478469 | 3 | 0-27877 | EPI_ISL_481739 | 2 | 27877-29902 | EPI_ISL_421783 |
| EPI_ISL_439137 | 0 | 0-8916 | EPI_ISL_488783 | 1 | 8916-29902 | EPI_ISL_439093 |
| EPI_ISL_518936 | 2 | 0-28878 | EPI_ISL_519052 | 0 | 28878-29902 | EPI_ISL_441368 |
| EPI_ISL_520231 | 2 | 0-28878 | EPI_ISL_519294 | 0 | 28878-29902 | EPI_ISL_499667 |
| EPI_ISL_437199 | 0 | 0-3095 | EPI_ISL_509695 | 3 | 3095-29902 | EPI_ISL_475921 |
| EPI_ISL_521429 | 2 | 0-28878 | EPI_ISL_518927 | 0 | 28878-29902 | EPI_ISL_499667 |
| EPI_ISL_440589 | 3 | 0-27877 | EPI_ISL_508873 | 2 | 27877-29902 | EPI_ISL_492160 |
| EPI_ISL_476327 | 3 | 0-25563 | EPI_ISL_444534 | 2 | 25563-29902 | EPI_ISL_470615 |
| EPI_ISL_451185 | 0 | 0-14804 | EPI_ISL_482575 | 2 | 14804-29902 | EPI_ISL_437090 |
| EPI_ISL_518513 | 2 | 0-28878 | EPI_ISL_519046 | 0 | 28878-29902 | EPI_ISL_499667 |
| EPI_ISL_437299 | 0 | 0-3095 | EPI_ISL_513182 | 3 | 3095-29902 | EPI_ISL_475921 |
| EPI_ISL_519175 | 2 | 0-28878 | EPI_ISL_519657 | 0 | 28878-29902 | EPI_ISL_499667 |
| EPI_ISL_519782 | 2 | 0-28878 | EPI_ISL_519506 | 0 | 28878-29902 | EPI_ISL_438065 |
| EPI_ISL_513162 | 0 | 0-3095 | EPI_ISL_447669 | 3 | 3095-29902 | EPI_ISL_512934 |
| EPI_ISL_495226 | 2 | 0-18486 | EPI_ISL_471929 | 0 | 18486-29902 | EPI_ISL_454991 |
| EPI_ISL_495051 | 3 | 0-27877 | EPI_ISL_475033 | 2 | 27877-29902 | EPI_ISL_465859 |
| EPI_ISL_513007 | 1 | 0-20134 | EPI_ISL_417999 | 0 | 20134-29902 | EPI_ISL_437472 |
| EPI_ISL_519576 | 2 | 0-28878 | EPI_ISL_522228 | 0 | 28878-29902 | EPI_ISL_479960 |
| EPI_ISL_519550 | 2 | 0-28878 | EPI_ISL_519034 | 0 | 28878-29902 | EPI_ISL_438065 |
| EPI_ISL_520515 | 2 | 0-28878 | EPI_ISL_520316 | 0 | 28878-29902 | EPI_ISL_518855 |
| EPI_ISL_524226 | 0 | 0-3095 | EPI_ISL_486105 | 3 | 3095-29902 | EPI_ISL_524368 |
| EPI_ISL_441321 | 3 | 0-27877 | EPI_ISL_456055 | 2 | 27877-29902 | EPI_ISL_457804 |
| EPI_ISL_464547 | 0 | 0-27383 | EPI_ISL_464180 | 2 | 27383-29902 | EPI_ISL_455854 |
| EPI_ISL_420451 | 1 | 0-28835 | EPI_ISL_420449 | 2 | 28835-29902 | EPI_ISL_454454 |
| EPI_ISL_512995 | 0 | 0-23928 | EPI_ISL_447669 | 2 | 23928-29902 | EPI_ISL_507037 |
| EPI_ISL_452934 | 3 | 0-27877 | EPI_ISL_417336 | 2 | 27877-29902 | EPI_ISL_421199 |

***Supplementary Table 4.*** *List of SARS-CoV-2 genomes flagged by Bolotie as recombinant. For each unique GISAID identifier listed in column 1, we show its parental clades, approximate breakpoint position, and closest parents for each of the parental clades.*
